# A body map of somatic mutagenesis in morphologically normal human tissues

**DOI:** 10.1101/2020.11.30.403436

**Authors:** Ruoyan Li, Lin Di, Jie Li, Wenyi Fan, Yachen Liu, Wenjia Guo, Weiling Liu, Lu Liu, Qiong Li, Liping Chen, Yamei Chen, Chuanwang Miao, Hongjin Liu, Yuqian Wang, Yuling Ma, Deshu Xu, Dongxin Lin, Yanyi Huang, Jianbin Wang, Fan Bai, Chen Wu

## Abstract

Somatic mutations accumulated in normal tissues are associated with aging and disease. Here, we performed a comprehensive genomic analysis of 1,737 morphologically normal tissue biopsies (~ 600 cells each), mostly from the epithelia, of nine organs from five donors. We found that somatic mutation accumulations and clonal expansions are widespread, although with variable extent, in morphologically normal human tissues. Somatic copy number alterations were rarely detected, except for tissues from esophagus and cardia. Endogenous mutational processes like SBS1 and SBS5 are ubiquitous among normal tissues though exhibiting different relative activities. Exogenous mutational processes like SBS22 were found in different tissues from the same donor. We reconstructed the spatial somatic clonal architecture with sub-millimeter resolution. In epithelial tissues from esophagus and cardia, macroscopic somatic clones expanded to several millimeters were frequently seen, whereas in tissues from colon, rectum, and duodenum somatic clones were microscopic in size and evolved independently. Our study depicted a body map of somatic mutations and clonal expansions from the same individuals, and it revealed that the degree of somatic clonal expansion and enrichment of driver mutations are highly organ specific.

## Introduction

Somatic mutations occur naturally during cell division. Although most somatic mutations do not confer phenotypic changes, a life-long accumulation may eventually alter essential genes that regulate cell proliferation and death, thus contributing to somatic clonal expansions. This process is associated with many aging-related diseases^1,2^. Particularly, in human epithelium, a type of cells that is widespread in the human body and undergo frequent turnovers, the accumulation of specific somatic mutations known as “cancer drivers” can trigger malignant transformation and unrestrained cell growth. This underscores the importance of understanding how early somatic mutations and clonal expansions in morphologically normal human epithelium affects cancer initiation.

Recent studies have revealed the somatic mutation landscape of different normal human tissues, including the skin^3,4^, esophagus^5,6^, colon and rectum^7^, liver^8^, endometrial epithelium^9^, bronchia^10^, brain^11,12^, embryo^13^, urothelium^14,15^, and blood cells^16,17^, mostly through deep DNA sequencing of biopsied tissue samples. Other studies have implemented bioinformatic algorithms to detect somatic mutations from RNA sequencing data of normal human tissues^18,19^. While those studies have contributed greatly to our knowledge of mutation rates, driver genes, and mutagenic factors in normal tissues from different human organs, the tissue samples they analyzed usually came from different donors who have distinct germline backgrounds and life histories, thus making a cross-organ comparison challenging. Ideally, for such comparisons, we should study somatic mutations and clonal expansions in normal tissues collected from the same individual.

Here we combined laser-capture microdissection (LCM) and mini-bulk exome sequencing to systematically investigate the somatic mutagenesis in morphologically normal tissues collected from nine anatomic sites of autopsy samples from five donors. Our intra- and inter-individual analyses with high spatial resolution provide new insights into the organ-specific pattern of somatic mutation and clonal expansion, thus expanding our understanding of human aging and the early onset of tumorigenesis.

### Sequencing normal tissues from multiple organs of the same individual

To accurately assess the somatic mutagenesis in different human organs and to compare the mutation and clonal expansion profiles within and between individuals, we applied a consistent sampling strategy, utilizing LCM to sequence approximate 600 cells from each tissue biopsy. In five deceased organ donors (PN1, PN2, PN7, PN8, and PN9), we collected approximately 50 biopsies from each of the nine anatomic sites of autopsy samples (Fig. 1a, Supplementary Table 1), covering morphologically normal epithelia from bronchia, esophagus, cardia, stomach, duodenum, colon, rectum, and morphologically normal parenchyma from liver and pancreas (Extended Data Fig. 1). While assuring quality control, we subjected 1,762 biopsies to whole exome sequencing, excluding 25 samples with less than 10X average coverage depth from further analysis. The remaining 1,737 samples have an average sequencing depth of 56X (Fig. 1a, Supplementary Table 2). We performed somatic alteration identification and mutant clone analysis, using DNA from the peripheral blood cells of each donor as the germline comparators.

**Fig. 1.**
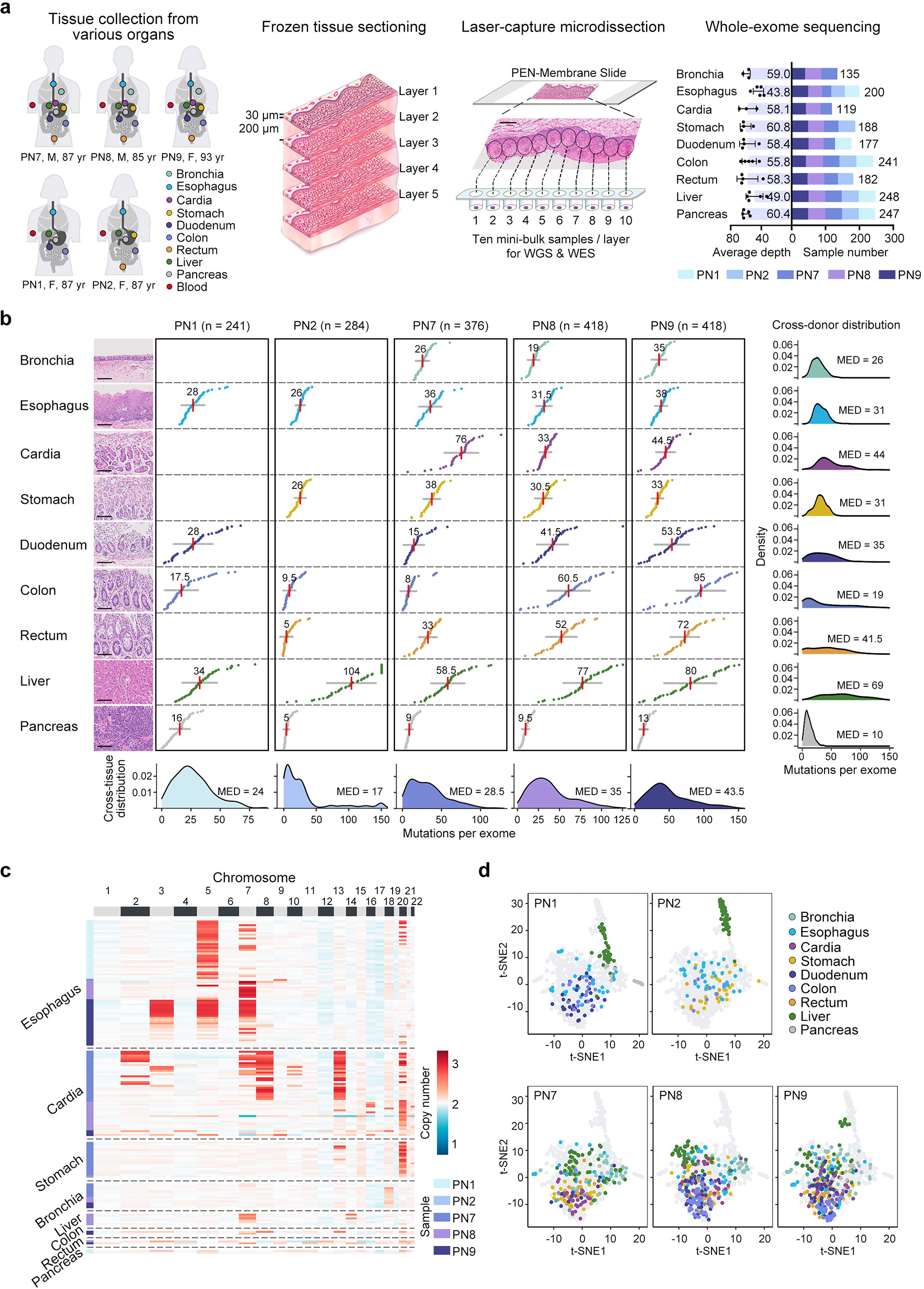
Research strategy and summary of somatic mutations detected in morphologically normal tissues of five donors (PNs). **a**, Laser-capture microdissections and mini-bulk exome sequencing procedures used to investigate somatic mutations and clonal expansions in nine anatomic sites of the autopsied samples. PEN, polyethylene naphthalate; WGS, whole genome sequencing; WES, whole exome sequencing. Scale bar = 400 μm. **b**, Numbers of somatic mutations detected in the coding regions in each tissue biopsy from the organs of five donors. Each biopsy sample is represented by a colored dot. Red vertical bars represent median mutation numbers and grey horizontal bars represent standard deviations. H&E stained tissues represent normal morphologies. Scale bar = 100 μm. **c**, Heat map showing somatic copy number alterations in biopsy samples. **d**, t-distributed stochastic neighbor embedding (tSNE) plots show the trinucleotide mutational spectra of biopsy samples from each donor. Only biopsy samples with more than 30 single nucleotide variations were included.

Overall, somatic mutations, both within and between individuals, displayed remarkable heterogeneity (Fig. 1b, Supplementary Table 3). Normal pancreas parenchyma tissue harbored the fewest number of mutations (median = 10/exome), consistent with the knowledge that the turnover of pancreas parenchyma cells is generally slower than that of epithelial cells. The mutational burden of normal liver tissues (median = 69/exome) was the highest among all normal tissues, substantially higher than those of epithelial cells from other organs. In particular, we observed the highest mutational burden (median = 104/exome) in the normal liver samples from PN2. In a cross-organ comparison, normal epithelial tissues from the cardia (median = 44/exome), rectum (median = 41.5/exome), and duodenum (median = 35/exome) harbored a higher number of mutations compared to those of the esophagus (median = 31/exome), stomach (median = 31/exome), bronchia (median = 26/exome), and colon (median = 19/exome). We also observed varying somatic mutation types and variant allele frequencies across all organs, which may reflect different underlying mutational processes and degree of mutant clonal expansions (Extended Data Figs. 2, 3).

### Somatic copy number alterations in normal tissues from different organs

We assessed somatic copy number alterations (CNAs) in normal tissues from different organs, by subjecting 1,764 biopsies to low-depth whole genome sequencing (Supplementary Table 4). Overall, we observed diploid genomes in most (1,608/1,764, 91.2%) normal tissue samples from different organs (Extended Data Fig. 4, Supplementary Table 4). However, sporadic CNA events can be detected in a number of samples (Fig. 1c). Interestingly, the samples with CNAs exhibited strong organ preferences. Normal esophagus tissues (31/41 [75.6%] from PN1, 10/41 [24.4%] from PN8, and 24/50 [48.0%] from PN9) were found to harbor CNAs that were enriched as whole chromosomal amplifications of chromosomes 3, 5, and 7. Yokoyama et al.^6^ reported an occasional amplification of chromosome 3 but not of chromosomes 5 and 7. A possible explanation is that our donors were significantly older than the study subjects enrolled in their study (87.8 vs. 63.7 year-old; P < 0.001; Wilcoxon rank-sum test). Also, in some cardia samples, (27/34 [79.4%] from PN7 and 15/44 [34.1%] from PN8), we detected CNA events that exhibited whole chromosomal amplifications of chromosomes 2, 7, 8, 13, and 20, which to the best of our knowledge has not previously been reported.

### Mutational signatures in normal tissues

We clustered the trinucleotide spectra of somatic mutations detected in normal tissue samples from different organs by using the t-distributed stochastic neighborhood embedding (t-SNE) projection (Fig. 1d, Extended Data Fig. 5a). Although most samples tended to cluster together independent of their origin of donors, some liver samples, mainly from PN1 and PN2, distributed separately from the main cluster.

To further explore the underlying mutational processes that operate in each of the normal tissues, we performed *de novo* single-base-substitution (SBS) mutational signature extraction based on a Bayesian hierarchical Dirichlet process^7,8^. To ensure enough mutation numbers for the analysis, we combined the mutations detected in LCM samples that we had collected within each slice (Supplementary Table 5). Totally, we deciphered seven mutational signatures (Sig. A to G), each of which mostly conformed to the Catalog of Somatic Mutations in Cancer (COSMIC) mutational signatures SBS5, SBS1, SBS22, SBS4, SBS45, SBS13, and SBS2, respectively^20,21^ (Fig. 2a, Extended Data Fig. 5b, c, Supplementary Tables 6, 7).

**Fig. 2.**
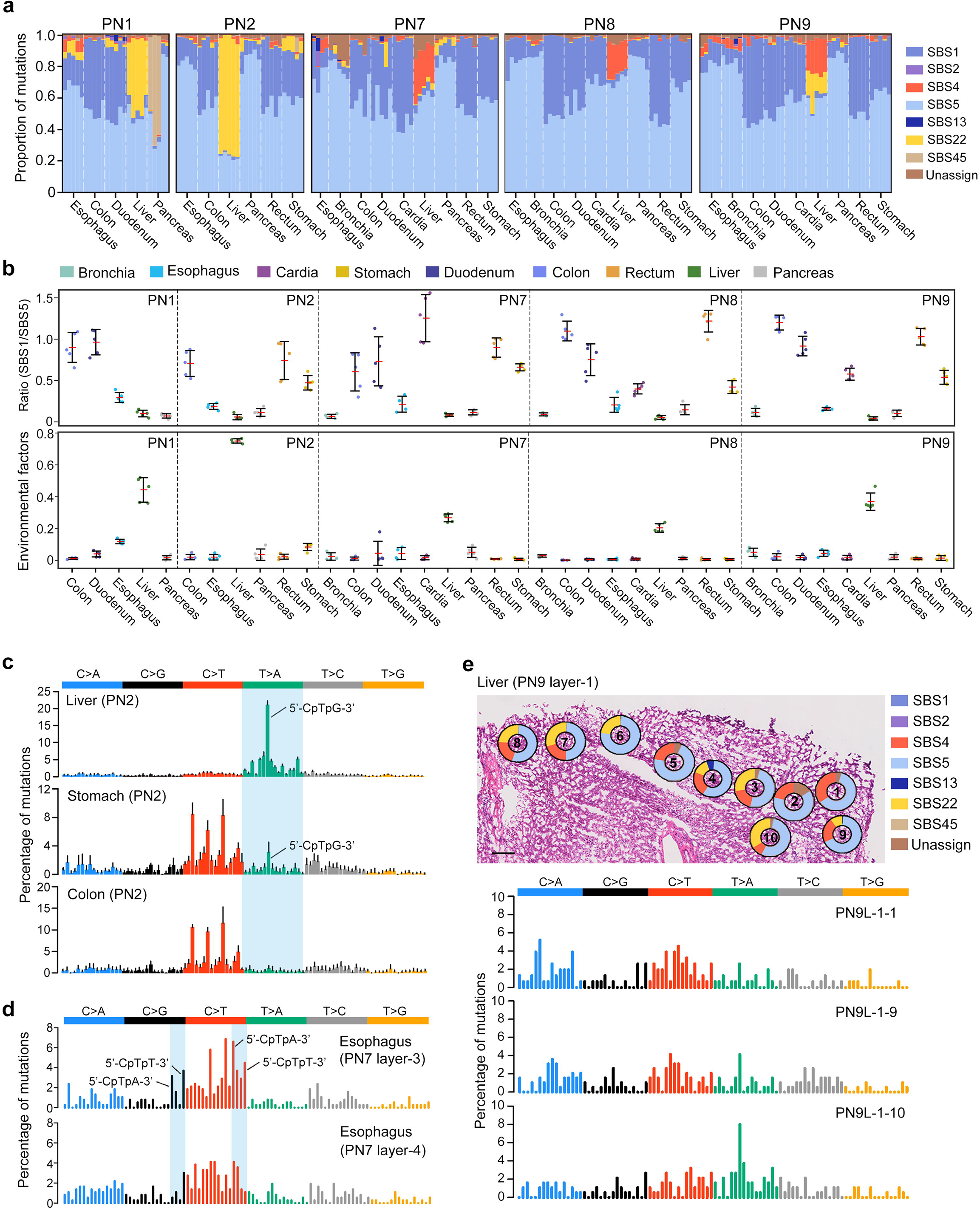
Mutational signatures of nine anatomic sites of autopsy samples from five donors. **a**, Stacked bar plots show the proportional contributions of each mutational signature in each organ of the five donors (PNs), as estimated through a Bayesian hierarchical Dirichlet process. Each stacked bar represents a combination of samples collected within each dissected layer. **b**, Upper: the proportion of SBS1 and SBS5 contributions across different organs. Lower: the summed contributions of SBS4 and SBS22 (Environmental factors) across different organs. Data are means (SD). **c**, Trinucleotide mutational spectra of liver, stomach, and colon tissues from PN2. The five dissected layers from each organ were combined. Each bar represents the mean (SD) of one of the 96 trinucleotide contexts. Typical AA-associated mutational features (shaded in blue) were obvious in liver and stomach, but absent in colon samples. **d**, Trinucleotide mutational spectra of two dissected layers of esophagus from PN7. Typical APOBEC-associated mutational features were shaded in blue. **e**, Representative example (PN9 liver) of the regional variations in mutational signature activities within a single dissected layer. Upper: representative H&E stained liver tissue with superimposed donut charts showing the proportional contributions of mutational signatures, as estimated by deconstructSig. Scale bar = 200 μm. Lower: trinucleotide mutational spectra of three representative laser capture microdissection biopsies.

We found two age-related endogenous mutational signatures, SBS1 and SBS5, throughout all normal samples from various organs and donors, suggesting the age-associated mutagenesis occurs ubiquitously in the human body (Fig. 2a, Extended Data Fig. 5d). Notably, we found that the relative contributions of SBS1 over SBS5 displayed a conserved organ-specific pattern across donors (Fig. 2b). The duodenum, colon, and rectum preferred to accumulate age-related mutations contributed by SBS1 rather than SBS5, while SBS5 showed relatively higher activities in the bronchia, pancreas, esophagus, and liver. The conspicuous preferences of these two clock-like mutational processes across organs may indicate some intrinsic tissue/cell-type biological differences, such as the cell proliferation rate. The preference of SBS1 over SBS5 activity has been compared among various cancer types^22^, but we investigated the preferences of somatic mutations detected in normal tissues from different organs. The two other endogenous mutational signatures, SBS2 and SBS13, emerged sporadically among normal samples (e.g., PN1 duodenum and PN7 esophagus; Fig. 2a) and are associated with the activity of APOBEC cytidine deaminases.

In addition, we identified two exogenous mutational signatures (SBS4 and SBS22), demonstrating that exposure to certain environmental stimuli also contributes to somatic mutagenesis in normal tissues (Fig. 2a, Extended Data Fig. 5d). We observed SBS4, the tobacco smoking associated mutational signature, in bronchia, esophagus and liver samples, consistent with previous findings^5,8,10^. Also, SBS22, whose underlying etiological factor is aristolochic acid (AA) exposure, exhibited notable activity in three donors’ liver samples (PN1, PN2, PN9) (Fig. 2a, Extended Data Fig. 5d). AA is a herb-extracted compound that has been extensively implicated in Asian liver and bladder cancers^23–25^. Of note, AA mutagenesis in alcohol-related liver disease and normal urothelium has recently been reported^8,15^. Intriguingly, the three donors with obvious SBS22 activity in our study were females (Supplementary Table 1), and such AA mutagenesis gender preference has been previously reported in upper tract urothelial carcinoma, but the underlying mechanism remains unclear^26^. We consistently observed that the normal liver tissues harbored more somatic mutations caused by exogeneous mutational processes (SBS4 or SBS22) than did the other organs, thus indicating a higher environmental carcinogen exposure risk for the liver than for other organs (Fig. 2b, Extended Data Fig. 5d).

Our sampling strategy largely enabled us to compare mutational signatures across different organs of the same donor, with the assumption that such samples were influenced by the same life history. The potential AA exposure history of PN1 and PN2 has been found to contribute to the somatic mutagenesis in liver. Within those donors, we also found AA mutagenesis in other organs, such as the esophagus and duodenum samples of PN1 and stomach of PN2 (Fig. 2a, c, Extended Data Fig. 6a). To our knowledge, there has been no previous reports of AA mutagenesis in either normal or cancerous stomach/duodenum tissues. Additionally, SBS22 has seldomly been reported in esophageal cancers and was absent in previous studies of normal esophagus tissues^5,6^.

Spatially correlated mini-bulk sequencing helped demonstrate that mutational signatures can be highly heterogeneous, even in the same organ from the same donor. For instance, we observed a marked difference in the mutational spectra between two tissue layers in donor PN7’s esophagus samples, with a noticeable APOBEC-associated mutational process in layer 3 but not in layer 4 (Fig. 2d). Similar intra-tissue heterogeneity (ITH) of mutational signatures was also exemplified by SBS22 in PN7 duodenum samples (Extended Data Fig. 6b). ITH of mutational processes was even identified within single tissue layers that we collected. For instance, both exogeneous mutational signatures SBS4 and SBS22 exhibited high activities in PN9 liver samples (Fig. 2a). When further exploring individual LCM biopsies in each tissue layer, we found remarkable differences in mutational spectra and relative activities of SBS4 and SBS22, even between adjacent LCM biopsies (Fig. 2e, Extended Data Fig. 6c). This regional variation (both between and within tissue layers) in the mutational signature activity may reflect regional activations of different endogenous and exogeneous mutagenic driving factors.

### Genomic landscape of driver mutations

Driver mutations confer competitive and selective advantages on cells during somatic clonal evolution. We identified 32 potential driver genes, including canonical cancer drivers such as *NOTCH1, TP53, ARID1A*, and *ERBB2* (Fig. 3a, Extended Data Fig. 7a, Supplementary Table 8). These 32 genes recapitulated signaling pathways that have been widely implicated in tumorigenesis (Extended Data Fig. 7b, c). Dysfunctions of those key pathways may promote normal cell fitness during early clonal evolution.

**Fig. 3.**
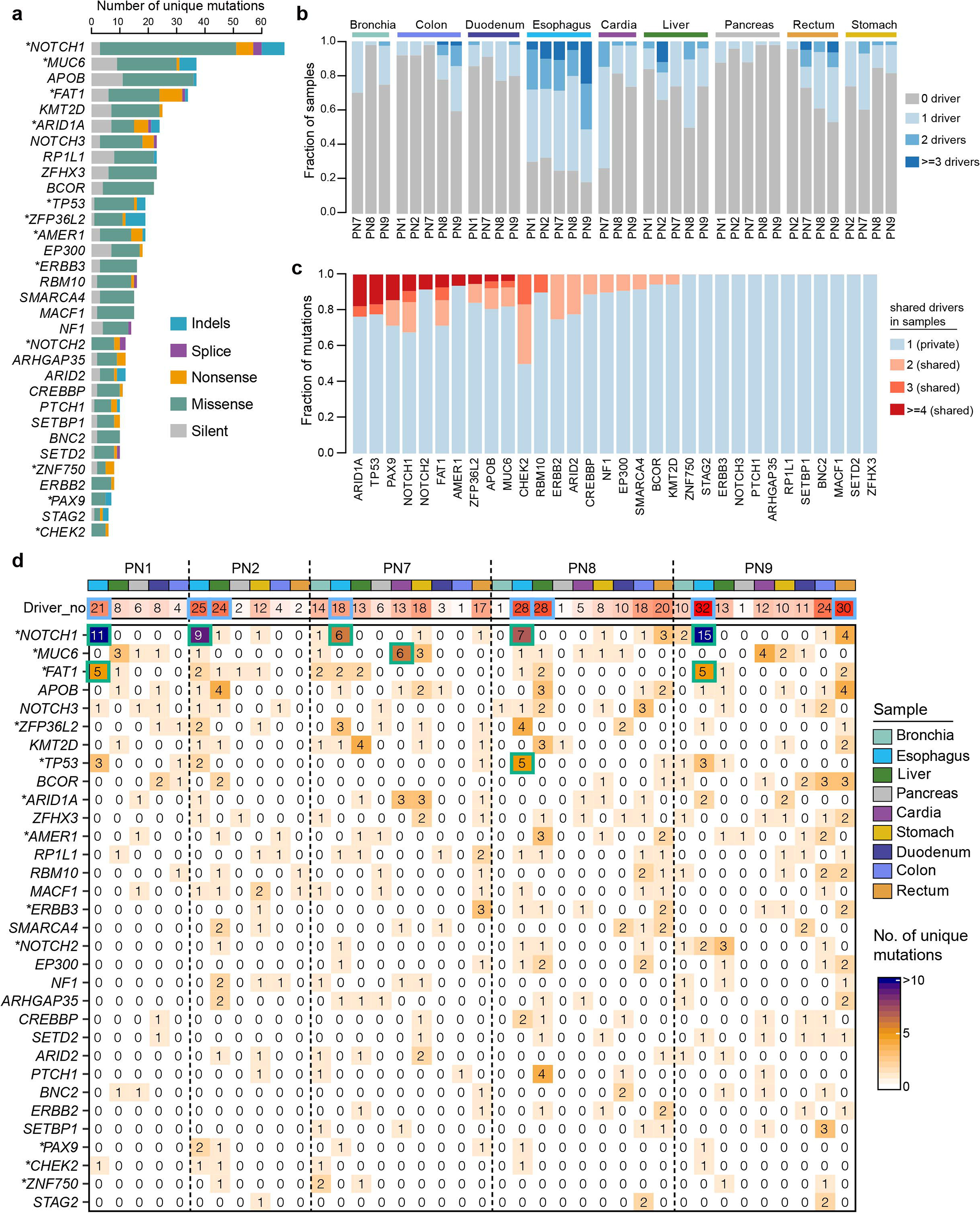
Mutational landscape of driver genes across organs. **a**, A stacked bar plot shows the number of unique mutations in each of the 32 driver genes. Asterisks indicate genes that were significant (q<0.1) in *dNdScv* analysis. **b**, Stacked bar plot showing the fraction of biopsy samples with zero, one, two, or three driver mutations across the five donors (PNs). **c**, Fraction of driver mutations that are private or shared by more than one biopsy samples. **d**, Heatmap showing the number of non-silent unique mutations in the 32 driver genes across different organs of the five donors. Blue boxes indicate organs with the highest number of driver mutations in each donor. Green boxes indicate representative driver genes with high mutation numbers in different organs and donors. Asterisks indicate genes that are significant (q<0.1) in *dNdScv* analysis.

The proportion of samples with driver mutations varied dramatically across organs. For instance, we detected driver mutations in about 6.5% of the pancreas parenchyma samples (ranging from 2% in PN9 to 12.2% in PN1) (Fig. 3b), but 73.8% of the esophageal samples harbored at least one driver mutation (ranging from 67.5% in PN2 to 81.6% in PN9) and about 11% of those samples (ranging from 4.6% in PN1 to 24.5% in PN9) harbored more than three driver mutations. On the other hand, we found mutations in genes such as *ARID1A, TP53, NOTCH1*, and *FAT1* tended to co-occur in multiple normal samples, which implied mutations in those genes were more likely to drive clonal expansions (Fig. 3c).

Mutations in the 32 potential driver genes distributed heterogeneously across both the nine organs and the five donors (Fig. 3d, Extended Data Fig. 7a). *NOTCH1* was found to be the most frequently mutated gene (65 unique non-silent mutations in 101 samples) (Fig. 3a, d). *NOTCH1* and *TP53* mutations, although widely observed across different organs, showed enrichments in esophageal tissues (R_o/e_: 2.87 and 2.92, respectively) (Fig. 3b, Extended Data Fig. 7d). While research had shown MUC6 to be a driver in gastric adenocarcinoma^27^, it was not reported as such in previous normal tissue sequencing studies. In our analysis, *MUC6* (28 unique non-silent mutations in 37 samples) was identified as a key driver gene that was frequently mutated in cardia and stomach tissues (R_o/e_: 6.59 and 2.14, respectively) (Fig. 3a, d, Extended Data Fig. 7d). Unexpectedly, the prevalence of *MUC6* mutations in normal gastric tissues (cardia and stomach) was significantly higher than that in gastric cancers (*P* < 10^−7^ using Fisher’s exact test; Extended Data Fig. 7e), suggesting that different molecular mechanisms may underlie clonal evolution of normal cells versus that of cancers. Similarly, mutated *AMER1* and *KMT2D* preferred liver tissues (R_o/e_: 2.21 and 2.76, respectively), *ERBB2* and *ERBB3* mutations occurred preferentially in rectal tissues (R_o/e_: 2.21 and 2.76, respectively), and *SMARCA4* and *CREBBP* mutations were enriched in duodenal tissues (R_o/e_: 6.12 and 4.9, respectively) (Fig. 3d, Extended Data Fig. 7d). We also observed heterogeneous driver mutation occurrences between individual donors. For example, we found four unique *PTCH1* mutations in PN8’s liver samples, but none in the other four donors’ liver samples (Fig. 3d).

### Spatial architecture of somatic mutant clones

We explored how somatic mutation accumulation and mutant clonal expansion coordinate in the normal tissues of different organs. For each donor, we plotted the distribution of mutational burden distribution versus the mutant cell fraction (MCF) of each sample (Fig. 4a, Extended Data Fig. 8). In esophageal and cardia tissues, mutant clones tended to be large, even if the mutational burdens were relatively low. In contrast, normal colon and rectal tissues accumulated many mutations, although the degree of clonal expansion was low. While in liver, some samples simultaneously harbored high mutational burdens and showed substantial clonal expansions. To further delineate how mutant clones evolved and expanded in normal tissues across organs, we combined phylogenetic analysis with mutation clustering and reconstructed the spatial clonal architecture with sub-millimeter resolution.

**Fig. 4.**
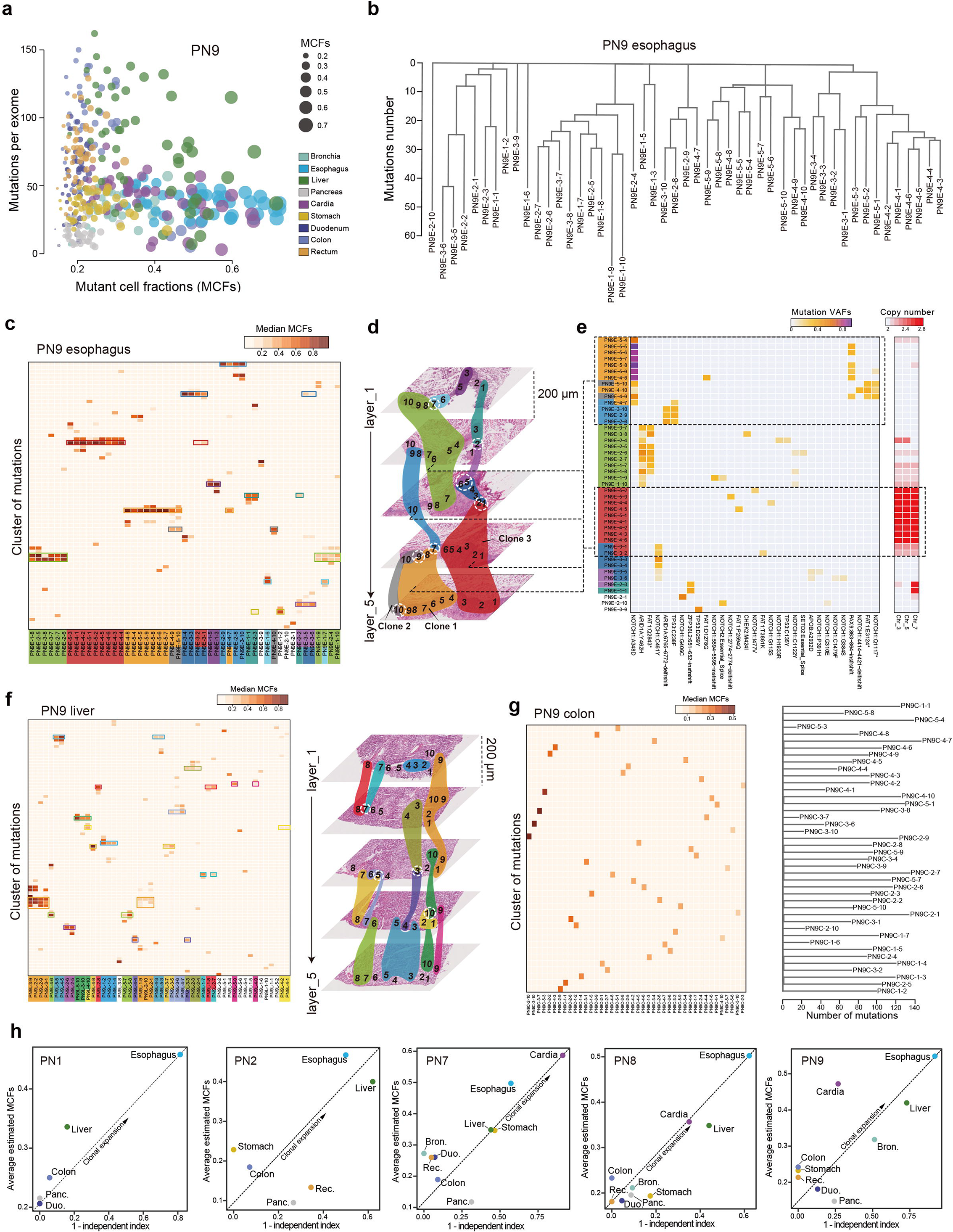
Estimation of somatic mutant clonal sizes and construction of spatial clonal expansion maps. **a**, Bubble plot showing the correlation between mutant cell fractions (MCFs) and the mutational burdens of biopsy samples across different organs in donor PN9. **b**, Phylogenetic tree depicting the clonal relationships of the biopsy samples in the esophagus of PN9. **c**, Heatmap showing mutation clustering in biopsy samples in the esophagus of PN9. Each cluster contains mutations with similar MCFs. The color scale represents the estimated median MCF for certain cluster in certain sample. Different colors shading the samples are used to indicate clonal sharing events among biopsy samples. **d**, Spatial clonal architecture of PN9’s esophagus tissues. The numbers in each layer represent the positions of laser-capture microdissection biopsy samples. The overlaid colors correspond to (**c**) and indicate the ranges of clonal expansions. Intermingling of different clones in single biopsy samples are highlighted using white dashed circles. **e**, Heatmaps showing potential driver mutations and copy number alterations in PN9’s esophagus samples. VAF, variant allele frequency. **f**, Left: Heatmap showing the mutation clustering in PN9’s liver samples. Right: Spatial clonal architecture of PN9’s liver tissue. **g**, Left: Heatmap showing the mutation clustering in PN9’s colon biopsy samples. Right: Phylogenetic tree depicting the clonal relationships of PN9’s colon biopsy samples. **h**, Correlations of average MCFs (estimated using a Bayesian Dirichlet process) with 1 minus independent index of the organs among the five donors.

We observed two major somatic clonal evolution scenarios across different organs in single donors: 1) A single mutant clone expands to a macroscopic scale, and 2) Competitive mutant clones originate and evolve independently. In PN9’s esophagus epithelium, we identified large-scale clonal expansions covering 2 to >10 LCM biopsies and spreading across two to three layers (Fig. 4b–d). Similar circumstances were also observed in other donors’ esophagus (Extended Data Fig. 9). Mutations such as those that occurred in *NOTCH1, TP53*, and *ARID1A* have potentially driven mutant clonal expansions in esophagus epithelium (Fig. 4e, Extended Data Fig. 9). For example, clone 1 harbored a *NOTCH1* mutation (p.A348D) and expanded to intermix with adjacent clone 2 which had a *FAT1* mutation (p.E3124*). We found that samples in clone 3 (a massively expanded clone spreading across three spatial dissected layers) shared no driver mutations. Interestingly, we found that samples in clone 3 ubiquitously carried CNAs in chromosomes 3, 5, and 7, which also recurred in esophagus samples from different donors in this study (Fig. 1c, Extended Data Fig. 9). This potential CNA-driven early clonal expansion in esophagus tissues has not previously been reported. The degree of mutant clonal expansion in liver tissue was comparable to that in esophagus tissues, but with fewer potential driver events being found (Fig. 4f, 3d). In a stark contrast, almost all colon and rectum samples we sequenced in PN9 were found to originate and evolve as independent mutant clones (Fig. 4g). These two typical scenarios were also observed in samples from other donors (Extended Data Figs. 9–11).

To profile the degrees of clonal expansions across different organs, we calculated an independent index and correlated the independent index with the average MCF in each organ. We defined a true clonal expansion as having a low independent index but a high average MCF. Being consistent across donors, we found that clonal expansions in esophagus, cardia, and liver were large while colon, rectum, and duodenum tended to have low degrees of clonal expansions (Fig. 4h). We also observed some inter-donor variation across donors. Mutant clones in the cardia and stomach tissues from PN7 were drastically expanded, which potentially associated with recurrent CNAs that they harbored (Fig. 1c, Extended Data Fig. 10). In PN2, the degree of clonal expansion in liver tissue was large and comparable to that in esophagus (Fig. 4h). This could have been related to the overwhelming AA mutagenesis in PN2’s liver tissues (Fig. 2a, Extended Data Fig. 5d).

## Discussion

In this work, we used laser-capture microdissections and mini-bulk exome sequencing to systematically characterize somatic mutation and clonal expansions in biopsy samples from nine anatomic sites collected from autopsy samples from five donors. Comparing tissue samples across the organs from the same individual could potentially offset potential systematic bias introduced by differing ages, germline backgrounds, or lifestyles when tissue samples from different individuals are compared. Our research strategy could reveal the intrinsic organ-specific patterns of somatic mutagenesis.

We observed stark intra-individual heterogeneity in the somatic mutational burdens, mutational signatures, driver landscapes, and degrees of clonal expansions in normal tissues from different organs. This is likely because of intrinsic differences in the physiological features and microenvironmental structures of each organ. Since the accumulation of somatic mutations is a direct result of cell division, it is natural to expect a positive correlation between mutational burdens and tissue turnover rates. However, since most cells in tissues are either partially or fully differentiated, and therefore typically short-lived, tissue turnover rates are difficult to estimate precisely. From a different perspective, a previous study estimated the total number of stem cell divisions in different tissues, and those numbers in colonic and rectal epithelia were noticeably higher than those of other tissues^28,29^. However, we confirmed higher somatic mutational burdens in the normal colonic and rectal epithelial tissues from PN8 and PN9, but not in PN1, PN2, and PN7. Our results suggest the correlation between somatic mutational burdens and total numbers of stem cell divisions in epithelial tissues might be complex.

Our within individual comparisons revealed an interesting pattern: the degree of somatic clonal expansion does not necessarily correlate with the mutational burden. Normal esophageal and cardia epithelial tissues showed more obvious clonal expansion compared to tissues from other organs. The greater somatic clonal expansion in esophageal tissue may be because those tissues harbor more driver mutations. It remains to be explored why somatic mutations in esophagus tend to occur in driver genes while the overall mutational burden is not high. Likewise, the obvious somatic clonal expansion in cardia samples remains mysterious. In PN8 and PN9, the normal cardia epithelial tissues were largely expanded, but in these samples, neither the total number of somatic mutations nor the number of driver mutations was high. In clear contrast, the colon and rectum samples from PN8 and PN9 harbored high mutational burdens, but the clonal expansion was weak. It has been previously suggested that somatic clonal expansion in colorectal tissues could be restricted to a single crypt with rare opportunity for further expansion^7,9^. Therefore, the differences between organs’ anatomic structures may be an important factor shaping somatic clonal expansions.

From the mutational signature analysis, we learned that somatic mutations and clonal expansions in normal tissues were mostly driven by the age-related mutational processes. On the other hand, the normal tissues we have studied here were constantly exposed to environmental stimuli (e.g., bronchia epithelia to air pollution and smoking, liver cells to metabolically active carcinogens). However, we found that the contributions from exogenous mutagenic factors were generally minor. Of note, the mutational signature associated with AA exposure, SBS22, was an exception, which was frequently seen in the normal liver samples and associated with a high somatic mutational burden. This highlights the notion that, in specific organs, somatic mutagenesis is driven more by environmental factors than by intrinsic factors. The dominant SBS22 we observed in liver samples may be a unique feature of Eastern Asians, which could explain the escalated incidence rate of liver cancer in Eastern Asia. It is intriguing that while SBS22 was observed in normal tissues of stomach, esophagus and duodenum in our study, it has not been reported in Chinese gastric cancer or esophageal cancer.

Compared to somatic mutations, CNAs are relatively rare and less extensive in the normal tissues from different organs, consistent with previous analyses^5,6,14,15^. In our study, liver samples harbored high mutational burdens coupled with obvious clonal expansions, however, CNAs were seldomly seen across the entire genome. In stark contrast, CNAs were frequently seen in normal esophageal and cardia samples. Why normal cells in those two sites are prone to harbor CNAs remains a mystery.

To summarize, our work provides a body map of somatic mutagenesis in morphologically normal human tissues. The organ-specific patterns of somatic mutation accumulation and clonal expansion aids our understanding of both aging and the early onset of tumorigenesis in various human organs.

## Acknowledgements

This project was jointly supported by the National Natural Science Foundation of China (81725015 to C.W., 81988101 to D.L. and C.W., 22050004 to J.W., 22050002 to Y.H.), Beijing Outstanding Young Scientist Program (BJJWZYJH01201910023027 to C.W.), Medical and Health Technology Innovation Project of Chinese Academy of Medical Sciences (2019-I2M-2-001 to D.L. and C.W.), National Science and Technology Major Project (2019YFC1315702, 2018ZX10302205 to F.B.), Guangdong Province Key Research and Development Program (2019B020226002 to F.B.), National Key R&D Program of China (2018YFA0108100 to Y.H.), Beijing Advanced Innovation Center for Genomics, and Shenzhen Bay Laboratory.

## Author Contributions

F.B., C.W., Y.H., J.W. and R.L. conceived the study; R.L., L.D., J.L., L.L. and D.X. performed data analyses; W.F., L.D. and J.L. performed experiments with assistance from Y.L., W.G., W.L., L.L., Q.L., L.C., Y.C., C.M., H.L., Y.W., Y.M. and D.X.; R.L., F.B., W.F., L.D. and J.L. wrote the manuscript with inputs from D.L., Y.H., J.W. and C.W.; Y.H., J.W., F.B. and C.W. supervised all aspects of this study.

## Materials and Methods

### Patients and sample collection

We obtained normal tissue samples from five deceased organ donors who had been recruited at the Body Donation Registration & Receiving Station in Peking Union Medical College, Beijing. Written informed consent was obtained from all donors. The study protocol and informed consent documents were reviewed and approved by the Local Research Ethics Committee (20/069-2265). None of the donors had undertaken neoadjuvant systemic therapy. Within 16 hours of the death, we separately collected tissues (lengths ranging from 1 to 5 cm) from nine organs (bronchus, esophagus, cardia, stomach, duodenum, colon, rectum, liver, and pancreas) from each donor. We then opened luminal organs longitudinally and cut each into approximately 0.5 × 0.5 cm pieces. All tissue samples were snap-frozen in liquid nitrogen and stored at −80 °C. Each donor’s clinical and pathological characteristics are summarized in Supplementary Table 1.

### Histopathological examination

We fixed the tissues in 10% buffered formalin and embedded them in paraffin blocks. Then the formalin-fixed paraffin-embedded (FFPE) tissues were sectioned into 3 μm-thick sections. The specimens were hematoxylin-eosin (H&E) stained and analyzed under light microscopy. Three pathologists independently examined the tissues’ morphological and histological features. The slides were imaged using Vectra Polaris Automated Quantitative Pathology Imaging System (Perkin Elmer).

### Tissue section preparations

We embedded tissue sections in optimal cutting temperature (OCT) medium (Thermo Fisher Scientific, Waltham, MA) at −25 °C. A total of 5 layers were cut at a thickness of 30 μm using a Leica cryotome, with a 232 μm gap between each layer. We transferred each section to a polyethylene naphthalate membrane slide (Thermo Fisher Scientific) and then incubated the slides in cresyl violet acetate for 1 minute and rinsed them twice in water. The remaining, unmounted tissues were used for immunohistochemistry and immunofluorescence analyses.

### Laser capture microdissection (LCM)

We used an LMD7000 Laser Microdissection Microscope (Leica Microsystems, Wetzlar, Germany) with 10× magnification and proper laser settings to microdissect the mounted and stained tissues from the previous section. Only epithelial layers with a target size of 0.06 mm2, which corresponded to about 600 cells, were microdissected. We placed each 600-cell microdissected isolate into an empty cap of a nuclease-free 0.2 ml Axygen PCR tube (Thermo Fisher Scientific). We took photomicrographs both before and after LCM.

### Whole genome library preparation and sequencing

We lysed the LCM samples using a low-temperature protocol with cold-active protease to reduce DNA-base oxidative deamination, thus eliminating artifacts in somatic mutation calling. Specifically, each biopsy sample was lysed in 8 μl customized lysis buffer [15 μg/μl native Bacillus licheniformis Protease (Creative Enzymes, Cat. No. NATE-0633), 30 mM Tris-HCl (pH 7.6, Rockland Immunochemicals, Gilbertsville, PA, Cat. No. MB-003), 10 mM NaCl (Ambion, Cat. No. AM9760G), 5 mM EDTA (Ambion, Cat. No. AM9260G), 0.4% Triton X-100 (Sigma, Cat. No. T9284)] at 6°C for 1 hour. The lysate DNA was further tagmented by 1 μl Tn5 transposome (Vazyme, TTE Mix V50 in Cat. No. TD501) into adaptor-flanked fragments in 20 μl 1 × tagmentation buffer [10 mM Tris-HCl, 7 mM MgCl_2_ (Ambion, Cat. No. AM9530G), 10% N,N-dimethylformamide (Sigma, Cat. No. D4551), 4 × Protease Inhibitor (Promega, Cat. No. G6521)]. After incubating the tagmentation reaction at 55°C for 1 hour, 0.8 μM sequencing index primer and Q5 high-fidelity 1x master mix (New England Biolabs, Ipswich, MA, Cat. No. M0492) were added to perform PCR amplification. The PCR procedure was 10 mins at 72°C for gap-filling; 30 secs at 98°C for pre-denaturation; 21 cycles of 15 secs at 98°C, 30 secs at 60°C, and 2 mins at 72°C for denaturation; and 5 min at 72°C for the last elongation. The purified product was quality checked and sequenced using the Nextseq 500, Hiseq 4000, or HiSeq XTen sequencers (Illumina, San Diego, CA).

### Whole exome library preparation and sequencing

The sequencing libraries were exome captured using the SureSelectXT Human All Exon V6 (for esophagus libraries) (Agilent, Santa Clara, CA, Cat. No. 5190-8864) or V7 (for libraries from other tissues) (Cat. No. 5191-4005) following the manufacturer’s guidelines. The products were quality checked and sequenced with Illumina HiSeq XTen sequencers (Illumina, USA), generating 2 × 150 bp paired-end reads.

### Copy number analysis based on whole genome sequencing

We performed low-depth whole-genome sequencing (about 1.5 × 10^6^ uniquely-mapped reads) of each sample. For the data analysis, we first used Cutadapt^30^ to trim adapters from the paired-end reads. Then, the clean reads were mapped to human reference genome hg19 (University of California, Santa Cruz) by using Bowtie2^31^ with default settings. PCR duplicates were marked using Picard MarkDuplicates (http://Picard.Sourceforge.net). Unique reads were then tabulated into non-overlapping dynamic bins (500kb resolution) across the genome. Lowess regression normalization was performed to reduce GC bias of bin counts. Copy number was called using R package DNAcopy^32^ with the circular binary segmentation (CBS) algorithm. Finally, we calculated median absolute pairwise differences (MAPD) to identify and filter out low quality samples (MAPD > 0.2).

### Single nucleotide variant (SNV) and insertion/deletion (INDEL) calling

Paired-end reads from the sequencer were aligned to the human reference genome hg19 using the Burrows–Wheeler Aligner (BWA) with default parameter settings^33^. The aligned BAM files were then sorted and merged (if needed) using Samtools 0.1.19^34^. To call SNVs and INDELs from the exome sequencing data, we first realigned the mapped reads using the Genome Analysis Toolkit^35^ (GATK 2.1–8) based on information of the dbSNP 135 (www.ncbi.nlm.nih.gov/projects/SNP). Then, Picard-tools 1.76 was used to fix mate pairs and mark PCR duplicates (http://Picard.Sourceforge.net). Next, the base quality recalibration was performed with GATK.

#### SNV calling

We used MuTect^36^ (v.1.1.4) to call the SNVs in each biopsy sample, with the genomic DNA of white blood cells (WBC) from each donor’s peripheral blood as the germline comparator. To ensure SNVs calling accuracy, we applied a series of filtering steps. (1) At least 10-fold coverage was required in the WBC samples bearing at most 1-fold mutated coverage (One-fold mutated coverage was allowed only when the total local coverage in the WBC was over 50-fold). (2) At least 10-fold total coverage was required in tissue biopsy samples with at least 3-fold mutation coverage. (3) The mutation allele frequency of each SNV was required to be greater than 5%. (4) The minimum value of the mutated alleles’ maximum mapping quality score was 20. (5) Variants both listed in the dbSNP database and (6) reported by the National Heart, Lung, and Blood Institute (NHLBI) Exome Sequencing Project (ESP) (http://evs.gs.washington.edu/EVS) were removed.

#### INDEL calling

We used the GATK Unified Genotyper to call INDELs with a series of filtering steps. (1) At least 10-fold coverage was required in the WBC samples without any mutated reads. (2) At least 10-fold total coverage in tissue biopsy samples and no less than three-fold mutation coverage was required to support each INDEL. (3) Variants both listed in the dbSNP database and (4) reported by the National Heart, Lung, and Blood Institute Exome Sequencing Project were removed. All INDELs that passed the filtering process were manually reviewed using Samtools ‘tview’ to further eliminate those that presented in poorly mapped reads. We used SnpEff v.3.0^37^ to annotate all SNVs and INDELs

### Mutational signature extraction using the Bayesian hierarchical Dirichlet process (HDP)

We studied the underlying mutational signatures that operated in normal tissues from different organs based on all SNVs detected in both coding and non-coding regions in the exome sequencing data. In order to minimize the bias in mutational signature analysis, we combined normal biopsy samples from each dissected layer of each organ into a single sample, thus increasing the number of mutations available for the mutational signature analysis. Only unique mutations in each dissected layer were included to avoid double counting of mutations. We excluded donor PN8’s bronchia layer 1 sample from the following mutational signature analysis because the mutation number was smaller than 40.

Single-base-substitution (SBS) mutational signatures were extracted using the Bayesian hierarchical Dirichlet process implemented in the HDP R package^38^ (https://github.com/nicolaroberts/hdp). Substitutions (including C>A, C>T, C>G, T>A, T>C, and T>G) and their trinucleotide sequence contexts were considered in this analysis. Also, we defined no prior mutational signatures in this HDP-based analysis. Detailed information about parameter settings are included: (1) Hyperparameters for the α clustering parameter (α and β) were set to 1. (2) The parameter ‘initcc’ was set to 40 so that the extraction was started with 40 data clusters. (3) The initial 10,000 iterations of the Gibbs sampler (parameter ‘burnin’) were discarded. (4) After that, we collected 50 posterior samples (parameter ‘n’) with an interval of 50 iterations (parameter ‘space’). (5) After each Gibbs sampling iteration, three iterations of concentration parameter sampling were performed (parameter ‘cpiter’).

Then, we compared our extracted mutational signatures to those that have been reported and published (COSMIC and Pan-cancer Analysis of Whole Genomes [PCAWG] catalogs) based on the cosine similarity (the formula described below). Extracted signatures with cosine similarity >0.85 compared to a known signature from either the COSMIC or PCAWG catalogue of signatures were considered as the known signature with the highest similarity.

Cosine similarities between extracted mutational signatures (A) and known ones (B) were calculated as follows:

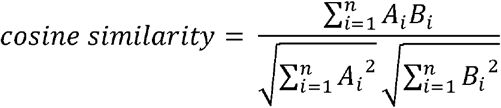

In this formula, n = 96 because we considered all 96 trinucleotide mutation contexts.

In total, we extracted seven mutational signatures which matched known mutational signatures. The relative activities of these signatures were used to generate the bar plot. The summed of SBS4 and SBS5 activities was considered as the mutagenic contribution from environmental factors.

### Mutational signature analysis using the deconstructSigs approach

We deconstructed the mutational signatures among donor PN9’s individual normal liver samples using the R package deconstructSigs^39^ v.1.8.0 with default parameters. This approach can identify the closest fit from linear combinations of pre-defined or known mutational signatures and can be used to decipher the relative activity of each signature in each sample using linear decomposition. In our analysis, we restricted the pre-defined mutational signatures to the seven that we *de novo* extracted (SBS1, SBS2, SBS4, SBS5, SBS13, SBS22, and SBS45) using HDP across normal tissues. We then used each signature’s relative weight to generate donut plots which were subsequently mapped onto histological photographs.

### Detection of potential driver genes under positive selection

To identify potential driver genes in the normal tissues of different organs, we used the dNdScv algorithm in R (https://github.com/im3sanger/dndscv)^40^, which calculates the ratio of the rate of non-silent mutations versus silent mutations, while considering the mutation sequence context, the sequence of each gene, and the mutation rate variation across genes. Since our entire study covered normal samples from nine different organs from five different donors, the driver detection process was complicated. Therefore, we adopted two strategies to detect diver genes using dNdScv. First, we included all somatic mutations detected in all normal tissues in the nine organs from the five donors as input for the dNdScv algorithm. Second, we included all somatic mutations detected in normal tissues in each specific organ from the five donors to do organ-specific dNdScv analysis. In each of those strategies, we used two gene sets (all coding genes and 126 selected driver gene candidates) for the hypothesis testing in dNdScv analysis. The 126-gene list included genes that selected from three sources: (1) driver genes that were identified using the dNdScv among cancers in the lung, esophagus, colorectum, liver, and stomach in a previous pan-cancer study^40^; (2) driver genes that were identified in The Cancer Genome Atlas lung^41^, esophageal^42^, colorectal^43^, liver^44^, and stomach^27^ cancer studies; (3) driver genes that have been reported in recent normal tissue sequencing studies regarding lung^10^, esophagus^5,6^, colorectum^7^, liver^8^, skin^3^, and endometrial^9^ tissues. The list of the 126 genes can be found in Supplementary Table 8. Through this analysis, we considered genes that matched one of the following five categories as potential driver genes in our study.

Category 1: Genes that were significant (q<0.1) when the analysis involved all tissues from all donors and hypothesis testing was applied across all coding genes.
Category 2: Genes that were significant (q<0.1) when the analysis involved all tissues from all donors and hypothesis testing was restricted to the 126 selected genes.
Category 3: Genes that were significant (q<0.1) when the analysis involved normal tissues in certain organs from the five donors and hypothesis testing applied across all coding genes.
Category 4: Genes that were significant (q<0.1) when the analysis involved normal tissues in certain organs from the five donors and hypothesis testing was restricted to the 126 selected genes.
Category 5: The genes that did not fit into any of the above four categories but harbored at least 8 unique somatic mutations in the 126 selected driver gene candidates.

We excluded *TTN* from the final list of potential driver genes because it mutates frequently in various cancers, most likely because of its large gene size. We conducted pathway enrichment analysis of our 32 putative driver genes using the Reactome FI Cytoscape plugin^45^.

To study the potential preference of *MUC6* mutations in normal cardia and stomach tissues, we listed the top-10 mostly frequently mutated genes in TCGA gastric cancer study and investigated number of mutations in these genes in normal cardia and stomach tissues. In TCGA gastric cancers, there were 13/266 mutations (in total of 266 mutations in these 10 genes) being detected in *MUC6*. While in normal cardia and stomach tissues, there were 17/30 mutations (in total of 30 mutations in these 10 genes) being detected in *MUC6*. We used the Fisher’s exact test to calculate the statistical significance.

### Mutated driver gene organ preferences

To explore the potential organ preferences of mutated driver genes, we compared the observed and expected number of mutations of each driver genes across different organs. The ratio of observation to expectation (R_O/E_) was calculated as follows:

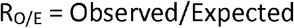

where the expected mutation numbers in different driver genes across organs were calculated based on the Chi-square test. We considered a driver gene both with R_O/E_ >1 in a specific organ and that was detected in that organ in more than one donor as having a potential preference for that organ.

### Mutant cell fractions and clone size

The fraction of cells with a somatic mutation in an individual normal biopsy sample is proportional to the variant allele frequency (VAF) of that mutation. However, that estimated fraction can be affected by local copy number alterations (CNAs) at the mutated site. As described in previous studies^3,5^, considering local copy number status, the relationship between the mutant cell fraction (MCF) and VAF of a specific somatic mutation can be described as:

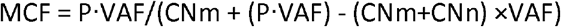

In this equation, P represents the average ploidy of cells without this mutation, CNm represents the copy number of alleles with the mutation in mutated cells, and CNn is the copy number of alleles without the mutation in mutated cells.

In the current study, we found most normal biopsy samples were free of CNAs. Meanwhile, we found only a small number of somatic mutations occurred in the genomic regions with CNVs in some specific normal tissues. Therefore, for mutations in the autosomes and × chromosome of females, the MCF can be simply calculated as follows: MCF = 2×VAF. While for mutations that occurred in the × chromosome of males, MCF can be estimated as follows: MCF = VAF. We only calculated MCFs of SNVs but not INDELs in our study.

### Clustering MCFs across multiple samples

To study the clone sharing events across multiple normal biopsy samples in different organs, we clustered somatic mutations from the multiple samples into different clusters based on the Bayesian Dirichlet process. We slightly modified the method described in a previous study^8^. Briefly, instead of using variant allele frequencies (VAFs) of mutations, we used MCFs of mutations (as integers) as input for the Bayesian Dirichlet process. As described in that study, the model includes a potential split-merge step at each cycle of the Gibbs sampler, followed by a previously described Metropolis–Hastings proposal for conjugate distributions. We ran the Gibbs sampler for 15,000 iterations, dropping the first 10,000 as a burn-in. We used the Equivalence Classes Representatives (ECR) algorithm^46^, implemented in the R package label.switching, to resolve the label-switching problem associated with mixture models. We removed clusters that harbored less than four somatic mutations.

### Defining the clonal independent index of each organ

We defined the clonal independent index of each organ in the five donors based on the results from the Bayesian Dirichlet process which was used to cluster MCFs of multiple biopsy samples. Briefly, the Bayesian Dirichlet process generates clusters of mutations and the estimated median MCFs of mutation clusters among biopsy samples in certain organ. We first ranked the estimated median MCFs and categorized them into 10 centiles. We considered that mutation clusters with estimated median MCFs larger than the second centile were located in a particular biopsy sample. Based on this, we calculated the number of biopsy samples that did not share any mutation clusters with others (independent sample number). Because we only considered mutation clusters with no less than four mutations as valid mutation clusters, the total number of biopsy samples in each organ in this analysis was calculated only considering those with at least one valid mutation cluster (total sample number). Finally, we defined independent index of an organ by dividing independent sample number by total sample number. We also calculated the average MCFs of the estimated median MCFs of valid mutation clusters. Finally, we considered that an elevated clonal expansion of mutant cells in a given organ was indicated simultaneously by high average MCFs and low independent index for that organ.

### Construction of phylogenetic trees and clonal expansion regions

We constructed phylogenetic trees in different organs to depict the clonal relationship of multiple normal biopsy samples using MEGAX^47^. Briefly, sequences with 3 base-pair in length surrounding all somatic mutations (including SNVs and INDELs) were extracted to construct the phylogenetic tree based on the maximum-parsimony algorithm. We regarded both SNVs and INDELs as single events and INDELs with different length contributed equally to SNVs in the construction of phylogenetic trees. Phylogenetic trees were further optimized by using Adobe Illustrator. Clonal expansion regions in esophagus and liver samples were according to the clonal sharing events revealed by clustering analysis regarding MCFs of mutations.

### Data availability

Whole-exome and whole-genome sequencing data generated in this study are being uploaded to the Genome Sequence Archive (GSA) in BIG Data Center (https://bigd.big.ac.cn), Beijing Institute of Genomics (BIG), Chinese Academy of Sciences, currently under the project number PRJCA003552.

### Code availability

Mutational signature analysis was performed using the hierarchical Dirichlet process R HDP package v.0.1.5 (https://github.com/nicolaroberts/hdp). Code for mutational signature analysis were adopted from https://github.com/HLee-Six/colon_microbiopsies. Code for the Bayesian Dirichlet process clustering of MCFs was adopted from https://github.com/sfbrunner/liver-pub-repo. Driver gene analysis was performed using the dNdScv v0.01 (https://github.com/im3sanger/dndscv).

---

**Extended Data Fig. 1.**
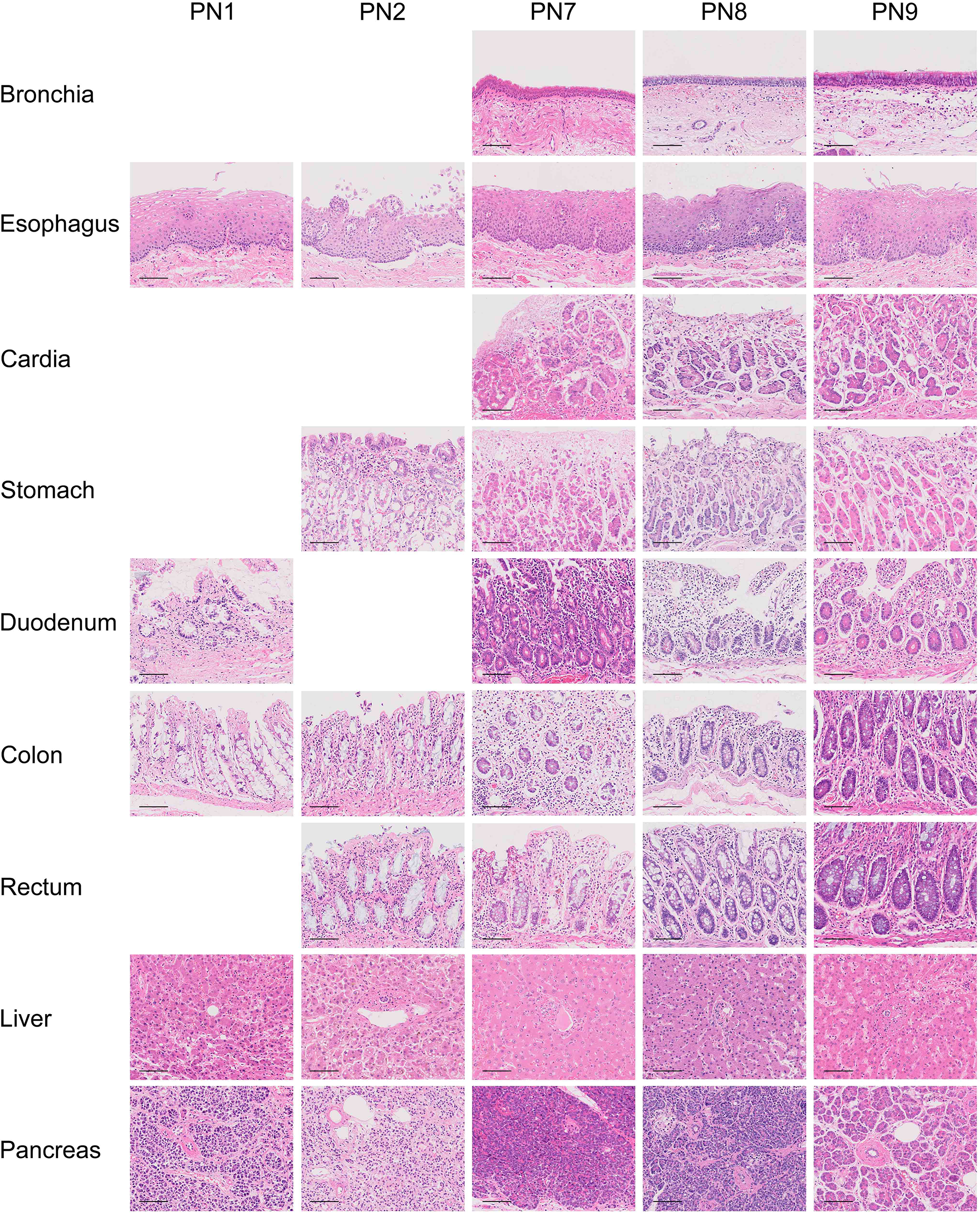
Normal tissue histology. Representative H&E stained samples showing the histological features of normal tissues sampled from nine organs from the five donors. Blanks in the figure means samples were not collected from corresponding organs and donors. Scale bar = 100 μm.

**Extended Data Fig. 2.**
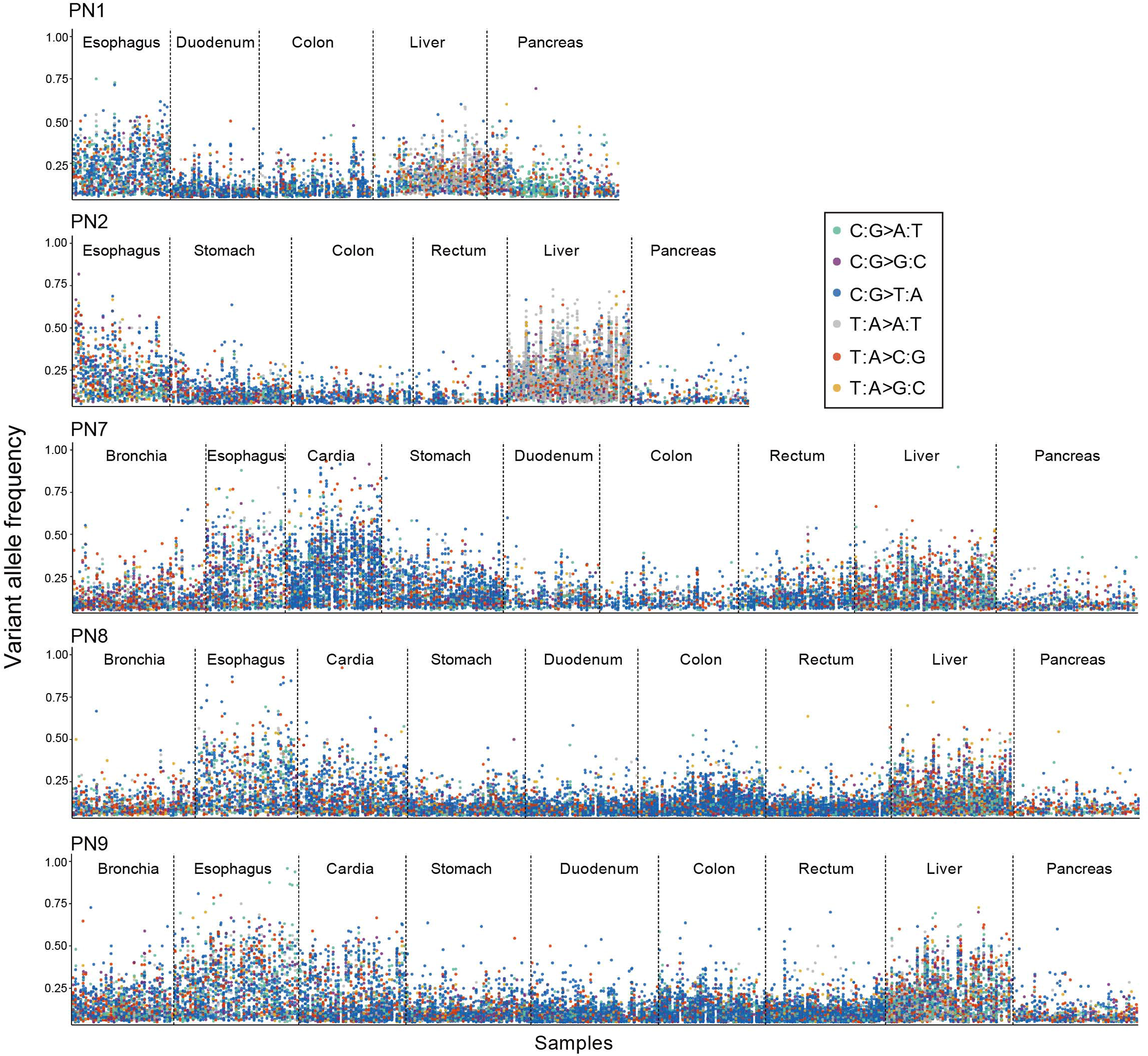
Variant allele frequencies (VAFs) of detected somatic mutations. Scatter plots showing the VAFs of somatic mutations detected in the normal tissues from the nine organs of the five donors (PNs). Dots are colored by mutation types.

**Extended Data Fig. 3.**
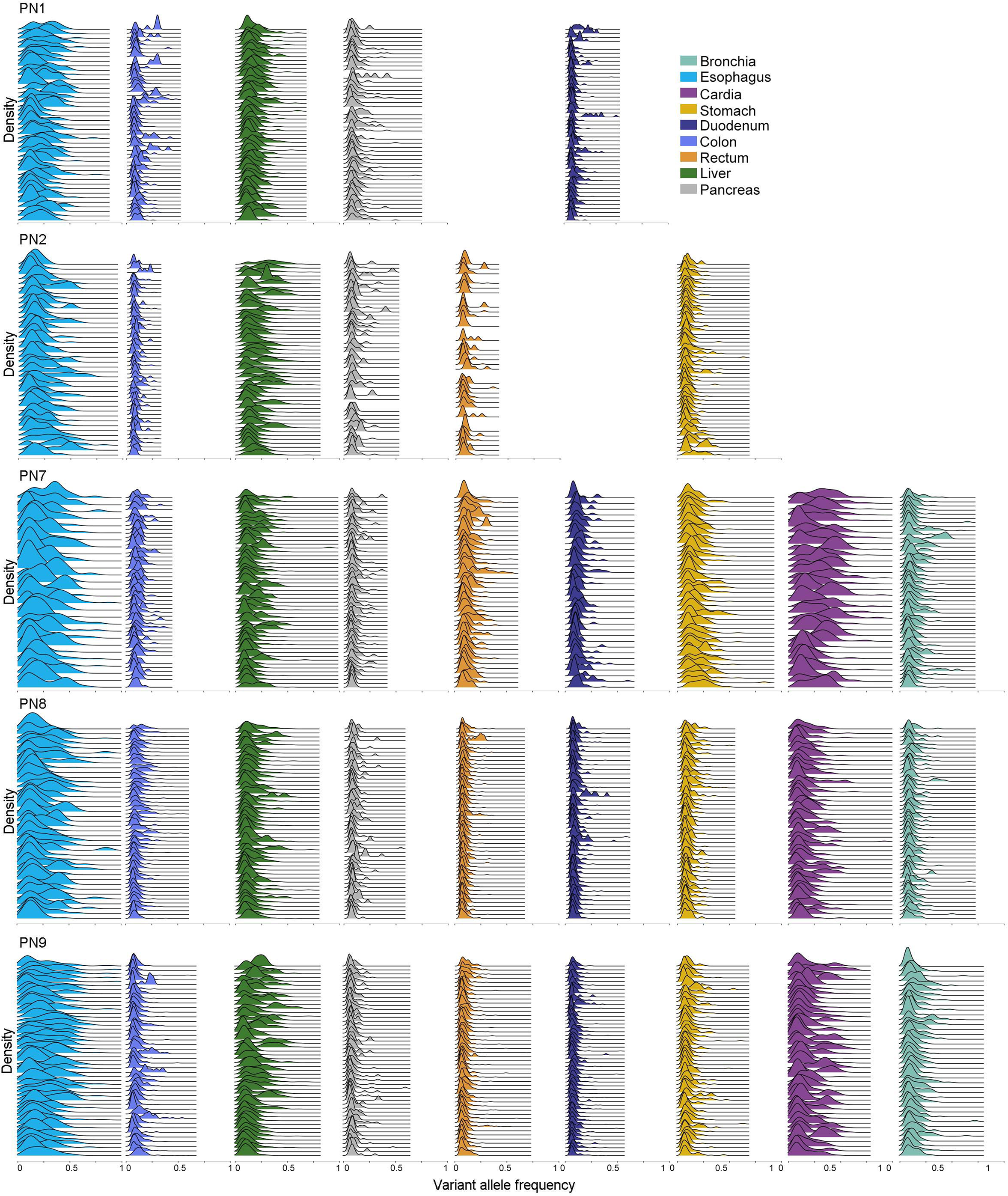
Density distributions of variant allele frequencies (VAFs). Ridgeline plots showing the VAF density distributions of somatic mutations detected in the normal tissues from the nine organs of the five donors (PNs) in this study, colored by organ types.

**Extended Data Fig. 4.**
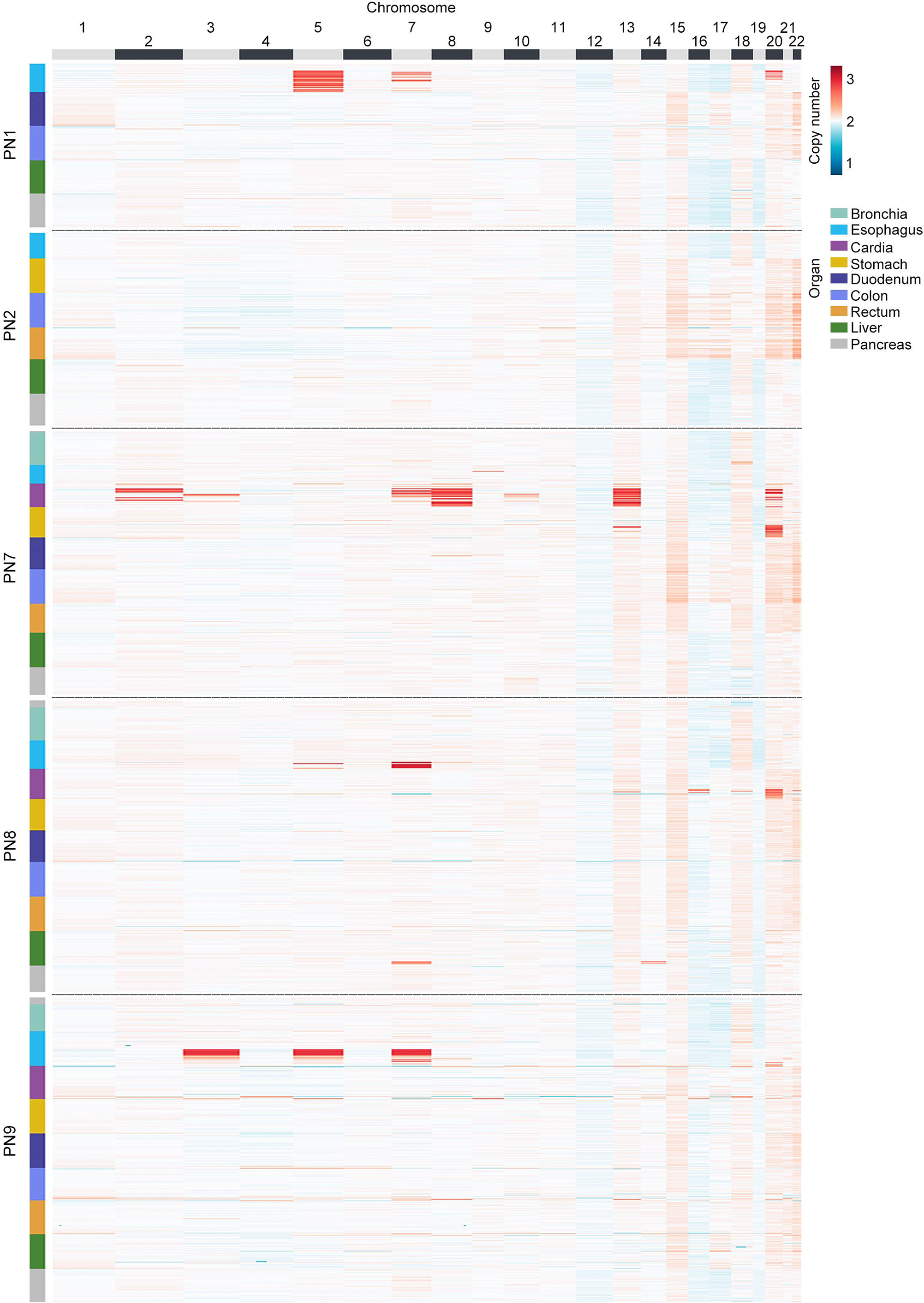
Somatic copy number alterations (CNAs). Heatmaps showing somatic CNAs detected in the normal tissues from the nine organs of the five donors (PNs). Sex chromosomes were excluded

**Extended Data Fig. 5.**
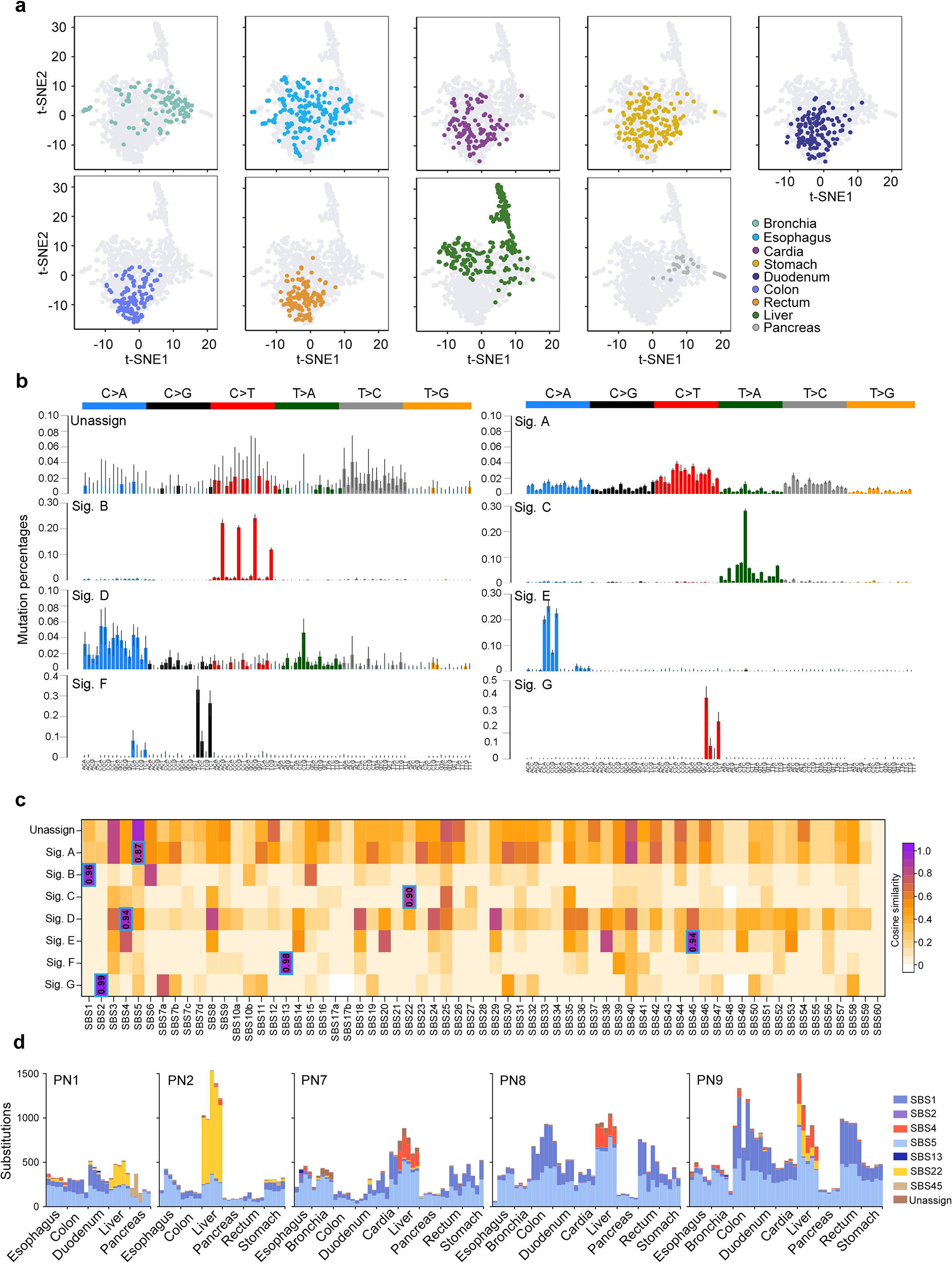
Mutational signature analysis. **a**, t-stochastic neighbor embedding (tSNE) plots of the trinucleotide mutational spectra of biopsy samples from each donor, broken down by different organs. Only biopsy samples with more than 30 single nucleotide variations were included. **b**, Trinucleotide mutational spectra for the unassigned signature and the seven signatures extracted using a Bayesian hierarchical Dirichlet process. The bars represent means (95% credible intervals) of the 96 trinucleotide contexts. **c**, Heatmap depicting the cosine similarities between extracted mutational signatures and mutational signatures from COSMIC and PCAWG catalogs. Cosine similarities between the seven extracted mutational signatures and their most similar comparators are highlighted. **d**, Stacked bar plots showing the number of mutations that caused by different mutational signatures.

**Extended Data Fig. 6.**
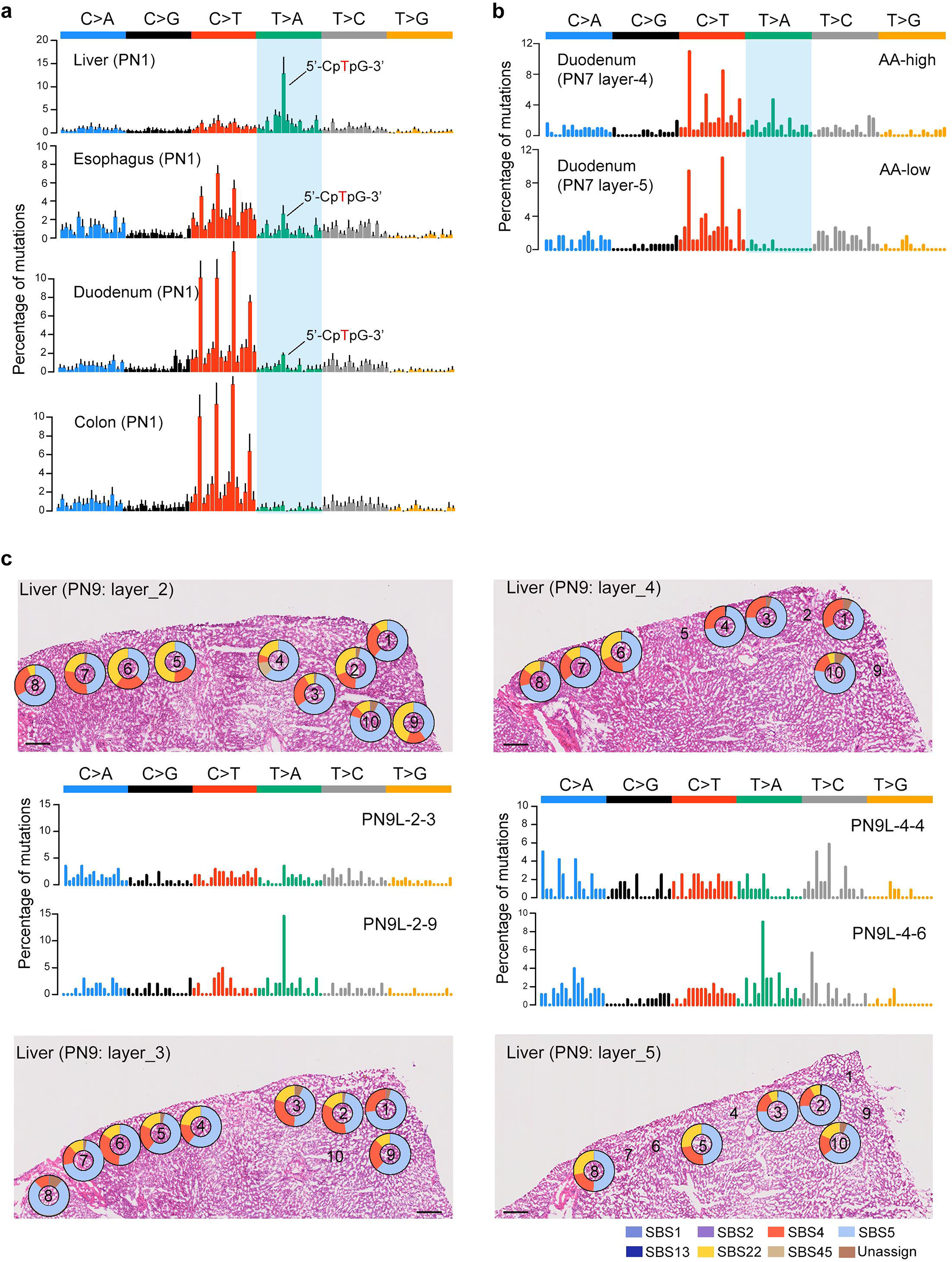
Intra-donor comparisons of mutational signatures. **a**, Trinucleotide mutational spectra of liver, esophagus, duodenum, and colon from donor PN1. Data from the five dissected layers of each organ were combined. Bars represent means (SD) of the 96 trinucleotide contexts. Typical AA-associated mutational features (shaded in blue) were obvious in liver, esophagus, and duodenum tissues, but absent in colon tissues. **b**, Trinucleotide mutational spectra of two dissected duodenum layers from PN7. Typical AA-associated mutational features (shaded in blue) were present in layer 4 but absent in layer 5. **c**, Representative example (PN9 liver) of regional variation in mutational signature activity within single dissected layers. Donut charts superimposed on representative H&E-stained tissues show the proportional contributions of mutational signatures estimated by deconstructSig. Scale bar = 200 μm

**Extended Data Fig. 7.**
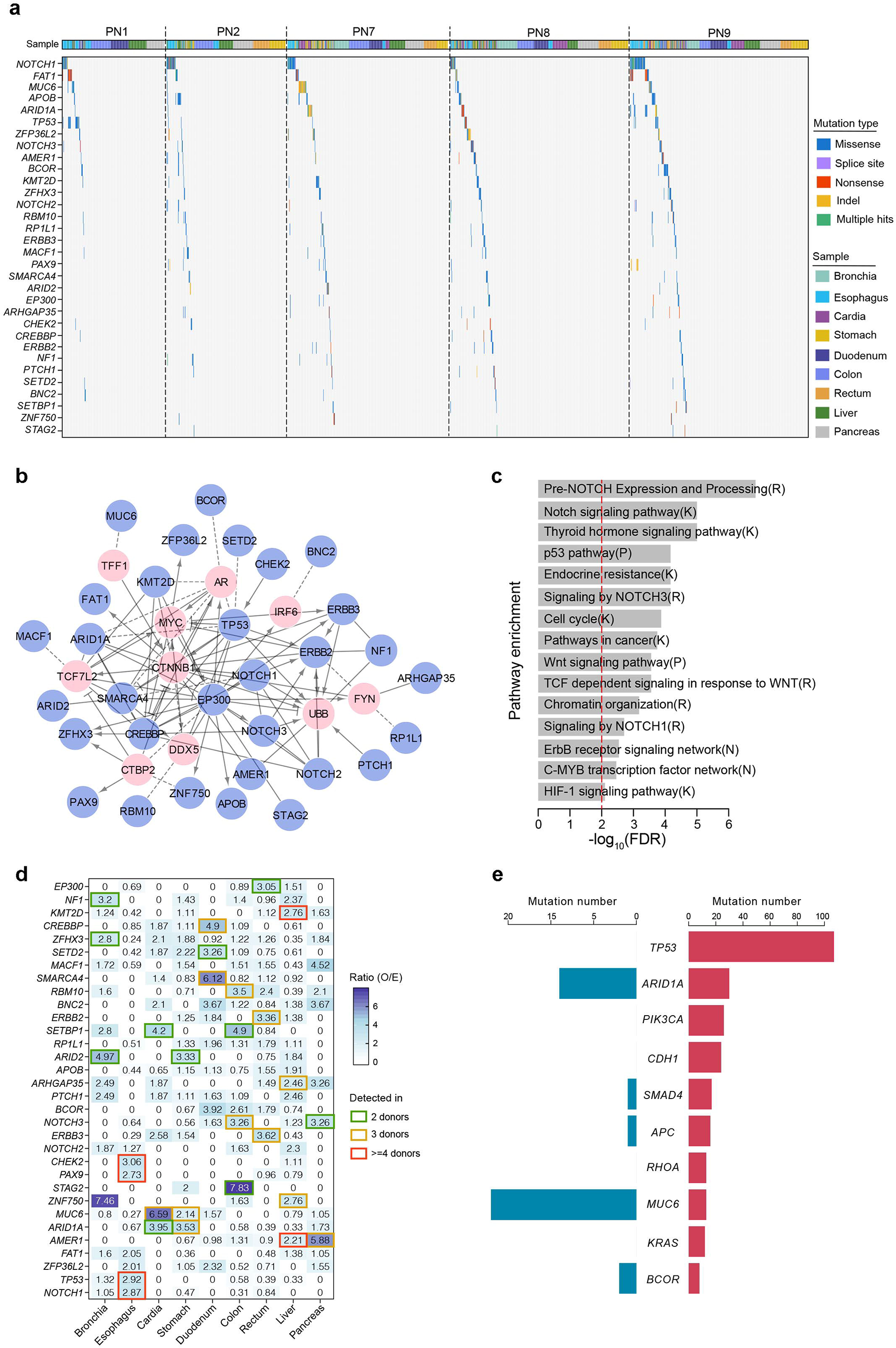
Landscape of driver mutations. **a**, Mutational landscape of the 32 putative driver genes across different organs from the five donors (PNs). **b**, The functional interaction (FI) network of the 32 driver genes. Driver genes are in blue nodes while linker genes (those not significantly mutated but highly connected to driver genes in the network) are in pink nodes. **c**, Significantly enriched pathways of the 32 driver genes. The vertical red line marks that false discovery rate (FDR) equals 0.01. **d**, Heatmap showing the ratio of the numbers of observed to expected (O/E) driver mutations across different organs. **e**, Bar plots comparing the number of mutations in gastric cancer top-10 most frequently mutated driver genes in The Cancer Genome Atlas (TCGA) with normal stomach and cardia samples in this study.

**Extended Data Fig. 8.**
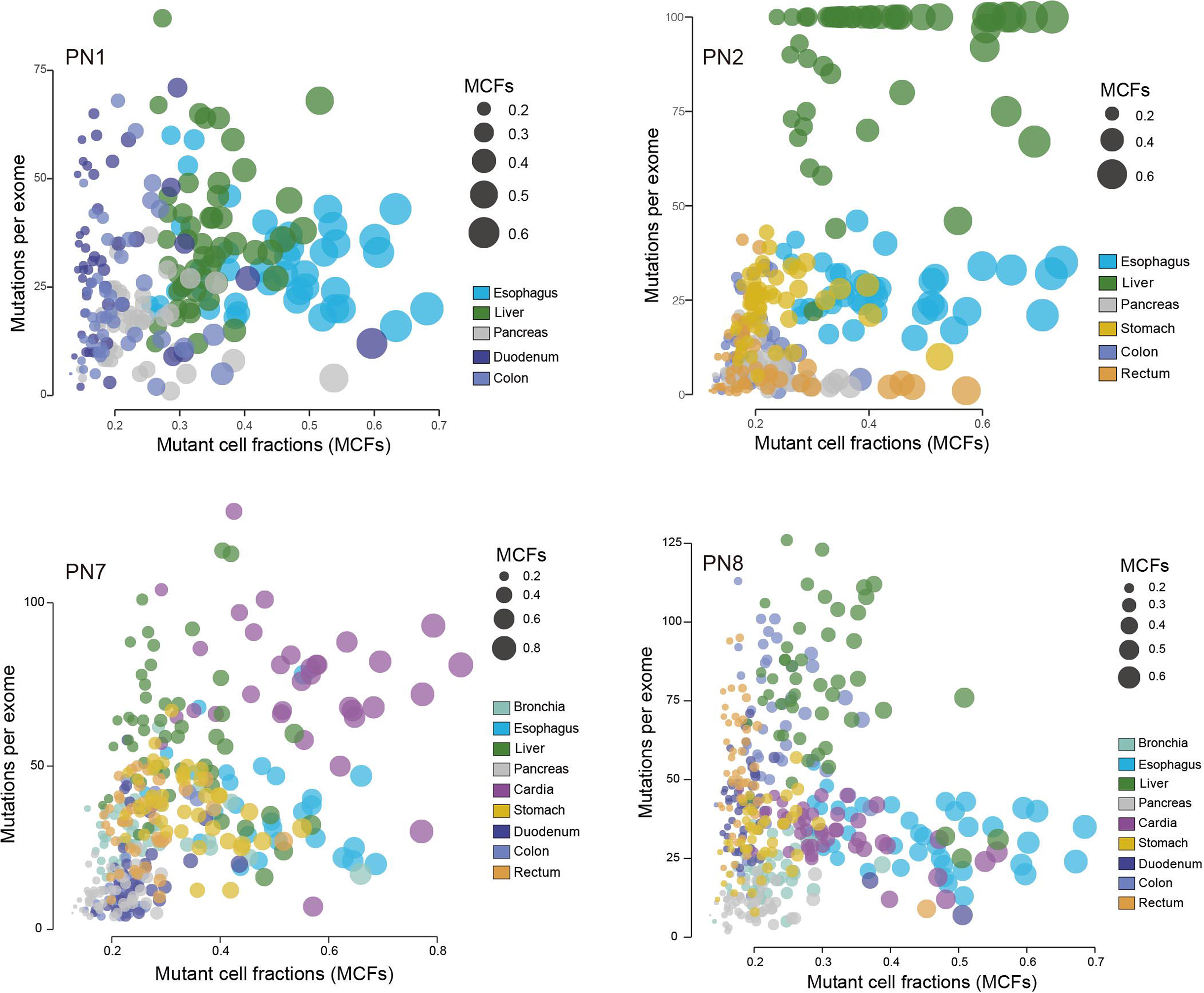
Relationships between mutational burdens and mutant cell fractions (MCFs). Bubble plots show the correlations between MCFs and mutational burdens in biopsy samples across different organs in donors PN1, PN2, PN7, and PN8.

**Extended Data Fig. 9.**
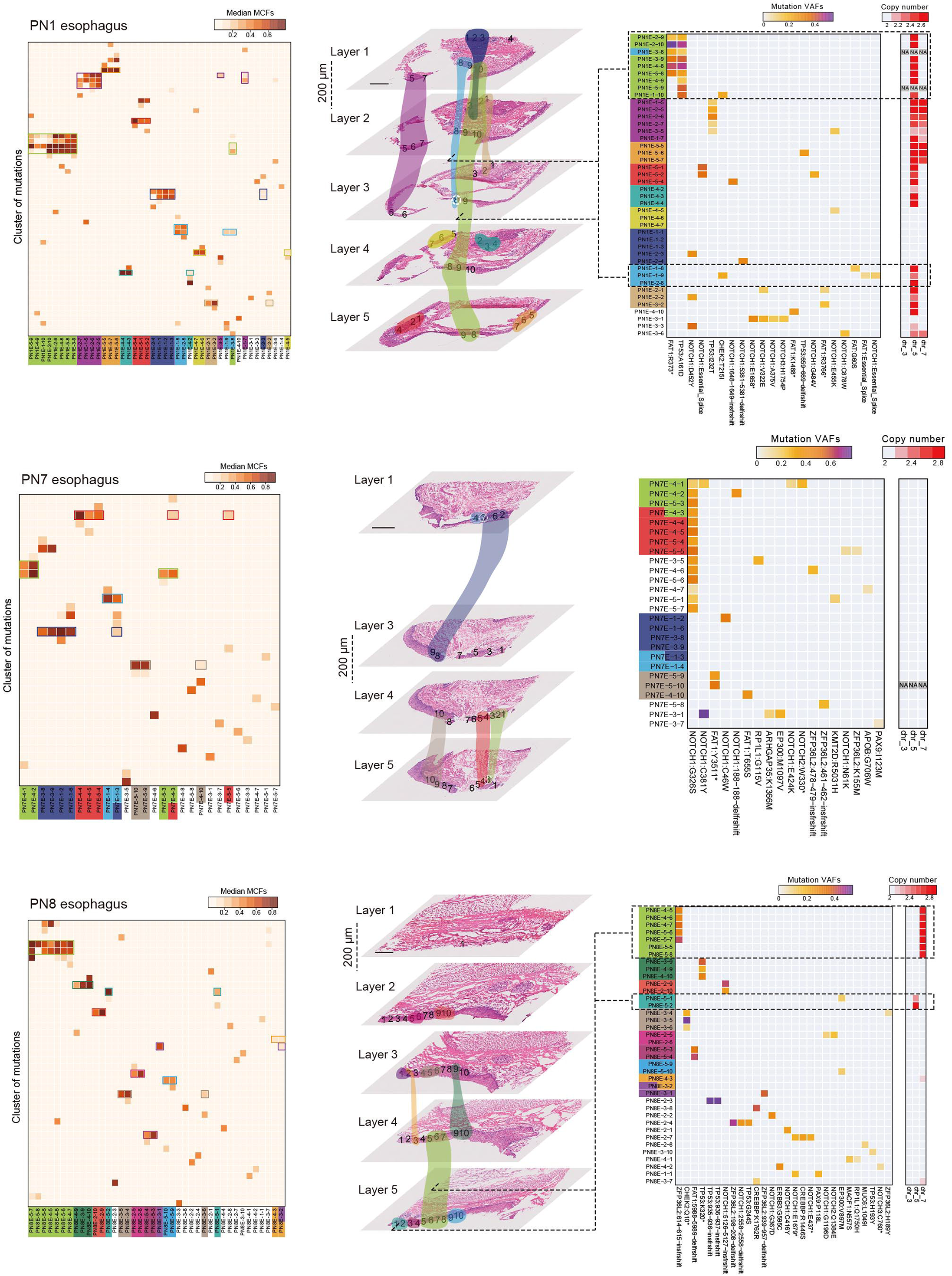
Mutant clonal expansion in esophageal epithelium. Heatmaps show mutation clustering, spatial clonal architecture, and potential driver mutations/CNAs in samples from esophagus. Each cluster contains mutations with similar mutant cell fractions (MCFs). Scale bar = 800 μm

**Extended Data Fig. 10.**
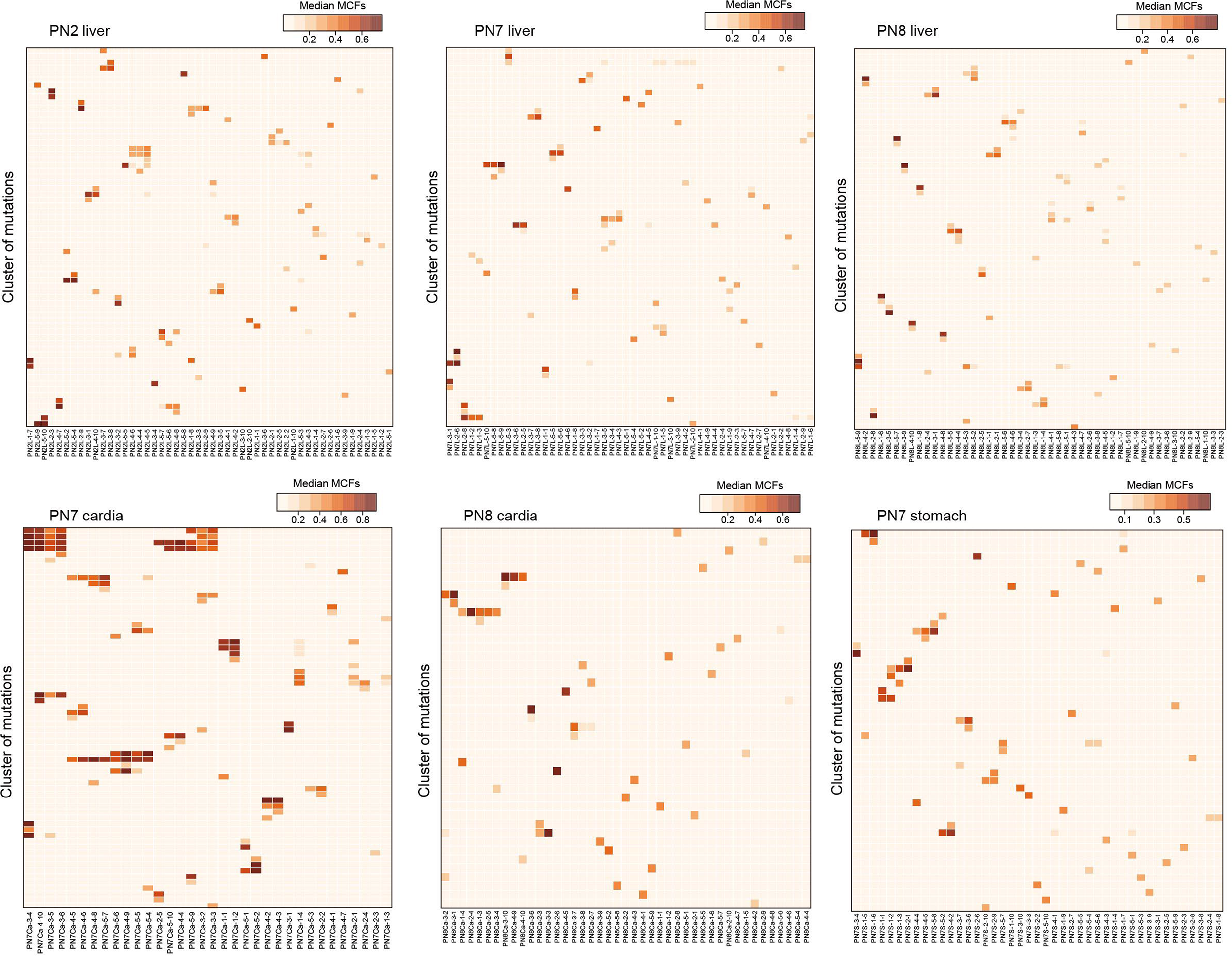
Representative examples of large scale mutant clonal expansion. Heatmaps show mutation clustering in samples from the representative organs. Each cluster contains mutations with similar mutant cell fractions (MCFs).

**Extended Data Fig. 11.**
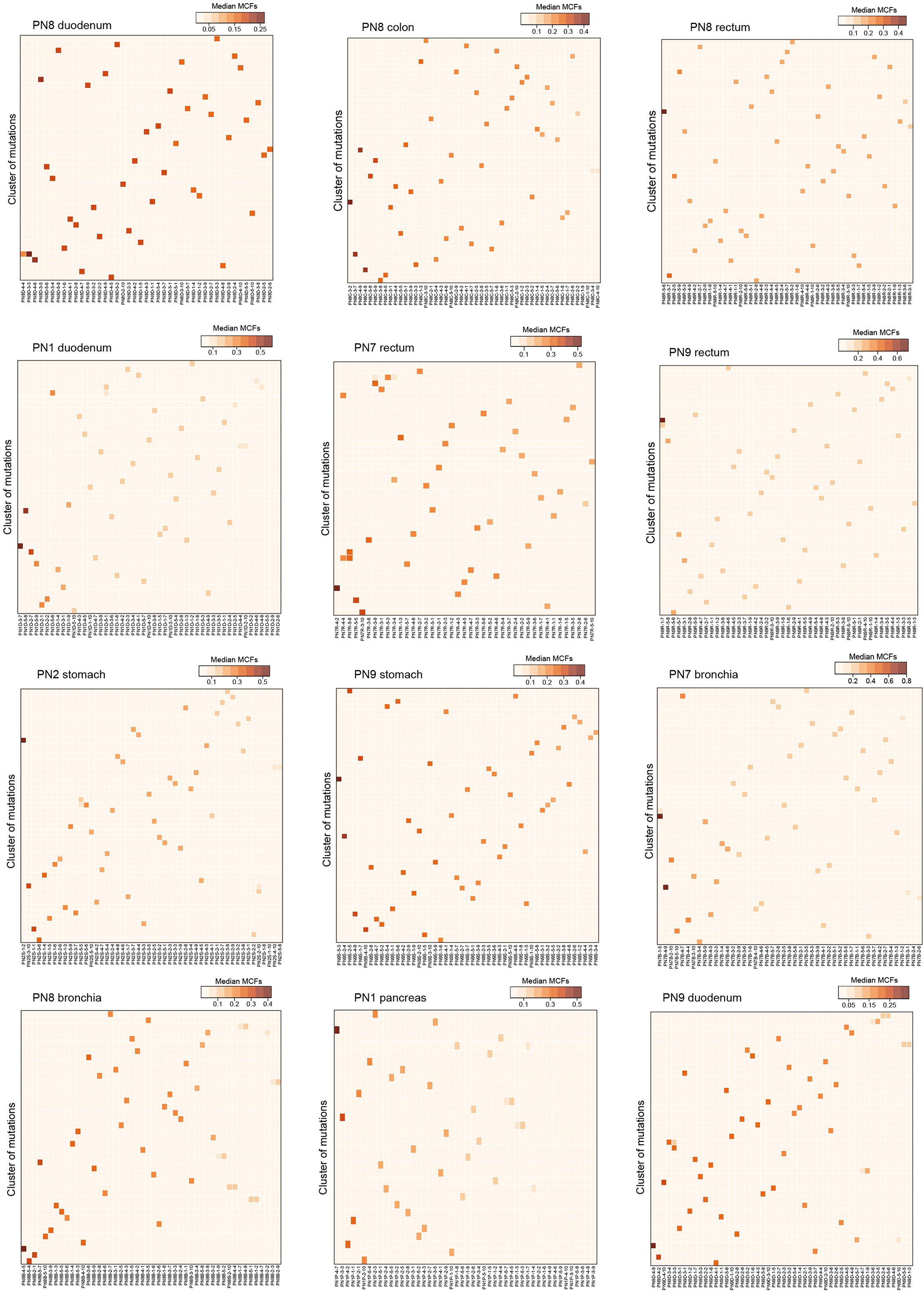
Representative examples of independent clonal evolution. Heatmaps showing clustered mutations in samples from representative organs. Each cluster contains mutations with similar mutant cell fractions (MCFs).

